# A high-quality, long-read *de novo* genome assembly to aid conservation of Hawaii’s last remaining crow species

**DOI:** 10.1101/349035

**Authors:** Jolene T. Sutton, Martin Helmkampf, Cynthia C. Steiner, M. Renee Bellinger, Jonas Korlach, Richard Hall, Primo Baybayan, Jill Muehling, Jenny Gu, Sarah Kingan, Bryce M. Masuda, Oliver Ryder

## Abstract

Genome-level data can provide researchers with unprecedented precision to examine the causes and genetic consequences of population declines, and to apply these results to conservation management. Here we present a high-quality, long-read, *de novo* genome assembly for one of the world’s most endangered bird species, the Alala. As the only remaining native crow species in Hawaii, the Alala survived solely in a captive breeding program from 2002 until 2016, at which point a long-term reintroduction program was initiated. The high-quality genome assembly was generated to lay the foundation for both comparative genomics studies, and the development of population-level genomic tools that will aid conservation and recovery efforts. We illustrate how the quality of this assembly places it amongst the very best avian genomes assembled to date, comparable to intensively studied model systems. We describe the genome architecture in terms of repetitive elements and runs of homozygosity, and we show that compared with more outbred species, the Alala genome is substantially more homozygous. We also provide annotations for a subset of immunity genes that are likely to be important for conservation applications, and we discuss how this genome is currently being used as a roadmap for downstream conservation applications.

## 1. Introduction

Whole-genome sequencing of threatened and endangered taxa enables conservation geneticists to transition from a reliance on limited numbers of genetic markers toward increased resolution of genome-wide genetic variation [*1,2*]. Such genome-level data offer unprecedented precision to examine the causes and genetic consequences of population declines, and to apply these results to conservation management [reviewed in *3,4*]. Moreover, continued decreases in the costs of genomic sequencing technologies make this information increasingly available for non-model organisms, including those with large genomes [*e.g. 5*; *6*]. Although challenges remain for bridging the gap between generating genomic data and applying this information to species management, this gap continues to close [for detailed discussions, see *3,7–10*].

Here we describe long-read, whole-genome sequencing and *de novo* assembly for the critically endangered Alala (Hawaiian crow; *Corvus hawaiiensis*). With 142 birds alive as of March 2018, this species is one of the most endangered avian species to have its genome assembled. The genome now provides valuable resources for conservation efforts, such as positional information and sequence data for candidate genes that are likely to have important fitness consequences (*e.g.* genes associated with immunity, mate choice, learning, and behavior). A high-quality genome also provides a tool for developing and mapping large numbers of genome-wide markers (*e.g.* single nucleotide polymorphisms [SNPs]), which can help to improve estimates of relatedness and individual inbreeding coefficients [*e.g. 11–14*]. Improved relatedness estimates will be important for choosing mating pairs in the conservation breeding (*i.e.* captive breeding) program, where inbreeding depression (*i.e.* loss of fitness due to inbreeding) has been observed during pedigree analysis [*15*]. The genome will also be used for comparative studies aimed at understanding the evolution of tool-use and other behaviors [*e.g. 16*]. Such comparative genome analyses could be especially important for conservation purposes, as they offer the potential to identify the genomic basis of traits associated with inbreeding depression both within and across species.

### 1.1. Study species and aims

Historically widespread within mesic and dry forest habitats on the Island of Hawaii, the Alala population declined rapidly during the twentieth century [*17*]. Fewer than 100 individuals were estimated to be alive in the 1970s, and the population continued to decline to fewer than 20 individuals during the 1990s. By 2002 the species was described as “extinct in the wild” by the International Union for Conservation of Nature and Natural Resources (IUCN) Red List of Threatened Species [reviewed in *18*]. Today, the Alala is one of the most endangered endemic bird species in Hawaii, having existed entirely in captivity from 2002-2016 [*15*]. All extant individuals are descended from nine genetic founders that established the conservation breeding program [described in *18,19*]. In 2016, a long-term reintroduction program was initiated in attempt to establish a self-sustaining population in the wild. Although a detailed pedigree has been established and utilized for captive management, including choosing breeding pairs, the current population exhibits signs of inbreeding depression [*15*]. For example, the species suffers from low hatching success, and the majority of inbreeding in the population appears to be attributed to a single founding pair [*18*]. Until establishment of the long-read genome assembly described here, molecular genetic studies were limited to small numbers of traditional genetic markers (*e.g.* microsatellite loci, amplified fragment length polymorphism [AFLP], and mitochondrial DNA markers [*20,21*]). These studies identified extremely low genetic diversity, which suggested that conservation efforts would benefit from a whole genome approach that could generate resources for targeting additional polymorphic regions (*e.g.* SNPs, and structural variation).

In this study we highlight the quality of the Alala genome assembly, and compare it to other avian assemblies that were also generated from whole-genome shotgun sequencing approaches. We provide details for a subset of candidate genes that we hypothesize will have important conservation implications, and we examine the repeat composition of the genome. We also describe analyses of runs of homozygosity (ROH) and the fraction of the genome estimated to be completely autozygous (fROH; [*22*]; *i.e.* identical by descent). Finally, we briefly discuss the goals and perceived challenges for the next stages of data generation and applications to Alala conservation and recovery.

## 2. Materials and Methods

### 2.1. Library construction and sequencing

Phenol-chloroform was used to extract high molecular weight genomic DNA from a blood sample taken from a single male Alala, named Hōike i ka pono (Figure S1, studbook # 32). This individual was chosen for genome sequencing because 1) his high inbreeding coefficient (0.25) would allow for simplified genome assembly, and 2) he is a great-grandson of the two genetic founders that constitute approximately 45% of the ancestry in the captive population (*i.e.* his genome would be a good representation for most birds in the population [*18*]). Library construction protocol followed the workflow for ultra large insert libraries [*23*]. The DNA was sheared to target 50 kb fragments (resulting distribution 30 – 80 kb) by using a Megaruptor^®^ (Diagenode), and assessed for quality by pulsed-field gel electrophoresis (PFGE) on the CHEF Mapper^®^ system (Bio-Rad). A total of 86 μg of DNA were then recovered from the 50 kb shearing condition. Sheared DNA was constructed into SMRTbell template by following the > 30 kb library construction protocol [*23*] with minor modifications (*e.g.* 1 × AM;Pure PB purification; room temperature rotation instead of vortexing; two-step elution process during AMPure PB elution to maximize recovery). Final SMRTbell library qualities were assessed by PFGE and Pippin Pulse (Sage Science) to determine the optimal size-selection cut-off of 20 kb. Size selection was done using the BluePippin system (Sage Science), with targeted exclusion of small fragments (< 20 kb) that would otherwise preferentially load during sequencing. Following size selection, the library fragments had a mode size of approximately 30 kb, and comprised approximately 8.6 μg of DNA; enough to sequence 133 single-molecule, real-time (SMRT) cells at Pacific Biosciences (PacBio). Sequence data were generated using the PacBio RSII instrument with P6v2 polymerase binding and C4 chemistry kits (P6-C4) and 6-hour run time movies, which yielded 128 Gb whole-genome sequence data. Post-filtering, the N50 subread length was 18,661 bp.

### 2.2. Genome assembly and quality

*De novo* assembly followed the PacBio string graph assembler process, using FALCON and FALCON-Unzip [*24*] to generate long-range phased haplotypes. During the assembly process, sequence reads were overlapped to form long consensus sequences [*6,24*]. These longer reads were used to generate a string graph, and the graph was reduced so that multiple edges formed by heterozygous structural variation were replaced to represent a single haplotype [*25*]. Primary contigs were formed by using the sequences of non-branching paths, while associated contigs (“haplotigs”) represent the sequences of branching paths. The resulting assembly thus represents a phased diploid genome [*24,26*].

To assess the quality of the final assembly, we compared the number and length of Alala contigs to those of other avian assemblies. In addition, we used BUSCO v2.0.1 [*27*] to assess the completeness of the gene space in the Alala assembly based on the detection of conserved single-copy orthologs. For comparison, we included genome assemblies of domestic chicken (*Gallus gallus,* GenBank accession GCF_000002315.4 [*28*]), Anna’s hummingbird (*Calypte anna*, GCA_002021895.1 [*26*]), zebra finch (*Taenopygia guttata*, GCF_000151805.1 [*29*]), hooded crow (*Corvus cornix cornix*, GCF_000738735.1 [*30*]), and American crow (*C. brachyrhynchos*, GCF_000691975.1 [*31*]) in the analysis. As lineage datasets, we chose eukaryota_odb9 (303 genes) and a 250-gene eukaryotic subset (pers. comm. Felipe Simão), which is highly congruent with the CEGMA dataset [*32*]. Gene finding parameters in the Augustus analysis step were based on the chicken genome.

### 2.3. Repeat composition analysis

To identify mobile and repetitive DNA in the Alala assembly, we generated a *de novo* repeat library using RepeatModeler v1.0.11 (repeatmasker.org). This software package primarily integrates RECON v1.08 [*33*] and RepeatScout v1.0.5 [*34*] to find interspersed repeats. Repeat models with 50% sequence identity over at least half their length to Swiss-Prot entries with known function were removed from the library, and remaining models were assigned to repeat classes by reference to Repbase (girinst.org). Additional, more detailed repeat classification was performed with CENSOR [*35*]. The Alala assembly was then screened for repetitive DNA using RepeatMasker v4.0.7 (repeatmasker.org) based on RMBlast and two repeat libraries: 1) the Alala repeat library described above, and 2) an expanded library also containing all chicken and ancestral eukaryotic repeats, as well as all zebra finch repeats provided by Repbase (accessed May 2, 2018). In addition to the version implemented in RepeatMasker, simple repeats were also assessed using the stand alone version of Tandem Repeats Finder v4.0.9 [*36*] with the following settings: Match = 2, Mismatching penalty = 7, Delta = 7, PM = 80, PI = 10, Minscore = 50, and MaxPeriod = 2,000. Along with the Alala assembly, we also analyzed the assemblies of the domestic chicken, Anna’s hummingbird zebra finch, hooded crow, and American crow listed above.

### 2.4. Candidate genes annotations

We focused on annotating particular genes associated with adaptive and innate immunity, as diversity at such genes is predicted to be especially relevant to fitness. Specifically, we were interested in genes of the major histocompatibility complex class II beta (MHC class II B), and toll-like receptor (TLR) genes. To identify candidate immunity genes in the Alala assembly, we first performed Blast (tblastn) searches using homologous protein sequences of other bird species as queries with an e-value cut-off of 1×10e^−5^. For MHC, queries were obtained from zebra finch, the currently best annotated passerine genome [*37*] (Table S1). For the more conserved TLRs, the full gene repertoire of the domestic chicken was used (Table S1). In the absence of transcriptional evidence, individual Alala genes were located by comparing genomic coordinates of high-scoring segment pairs on each contig, which often corresponded to exons. Genomic sequence including 1500 bp up- and downstream of each putative gene was extracted, and gene structure and coding sequence predicted by the AUGUSTUS web server v3.3 [*38*]. In the case of MHC class II B, only putative genes including exons 2 and 3 were considered for this step, due to a large number of single-exon or fragmentary hits. Portions of nucleotides that appeared to be missing from the predicted coding sequence were identified by aligning the predicted sequence to the reference using MAFFT v7 [*39*]. Using short sequence motifs taken from the reference, we then attempted to find missing homologous parts by translating the genomic sequence into all three reading frames in the coding direction by using EMBOSS transeq (http://www.ebi.ac.uk/Tools/st/emboss_transeq/). Finally, manually completed gene predictions were tentatively classified as functional, ambiguous or pseudogenized, depending on the integrity and length of the reading frame. Genes for which a complete reading frame including start and stop codons could be identified were considered functional, while genes that required the insertion or deletion of a single nucleotide to recover the complete reading frame (suggestive of a sequencing error) were categorized as ambiguous. Fragmentary reading frames or multiple frameshift mutations were regarded as indicative of pseudogenes. Untranslated 5’ and 3’ regions could not be annotated due to the lack of transcriptional evidence. MHC class II B Blast searches were later repeated to assess the number and divergence of gene fragments with increased sensitivity. Exons 2 and 3 of the Alala gene, Coha_MHCIIB_b (see Results), were used as query sequences.

To shed light on the evolutionary history of the gene family, we performed a phylogenetic analysis based on exon 2 of MHC class II B genes in Alala and other corvids. We included all functional and ambiguous Alala genes, as well as complete MHC class II B gene sets of single individuals each of American crow (13 genes), the jungle crow (*C. macrorhynchos japonensis*; 14 genes), the Asian rook (*C. frugilegus*; 11 genes) and the azure-winged magpie (*Cyanopica cyanus*, 7 genes) to allow for within-genome diversity comparisons. These data were obtained by Eimes *et al.* [*40* and pers. comm.) using a targeted PCR approach. Exon 2 nucleotide sequences were aligned with MAFFT v7, and 10 maximum likelihood trees computed under the GTRCAT model in RAxML v8.1.20 [*41*]. Confidence values were estimated from 500 rapid bootstrap replicates and drawn onto the best maximum likelihood tree (−f a algorithm).

### 2.5. MHC functional supertypes

To assess how similar the Alala MHC class II B repertoire might be to other corvids in terms of properties of the antigen-binding regions, we relied on comparisons of functional supertypes [*e.g. 42,43*]. Briefly, functional supertype analysis involves identifying codons under selection (positively-selected sites; PSS), and then grouping sequences according to descriptor variables that reflect the physical and chemical properties of the amino acids at these selected positions [*44–46*]. As the Alala MHC class II B sequences from this study were generated from a single individual, we based our analysis on the locations of nine PSS shared among three other crow species (jungle crow, carrion crow [*C. corone orientalis*], and American crow), which were previously identified through HYPHY analysis [*47*]. First, we used MUSCLE [*48,49*] implemented in Geneious vR10 [*50*] to align our putatively functional Alala exon 2 sequences to the 237 sequences from Eimes *et al.* [*47*], for a total of 244 nucleotide sequences. We then trimmed all sequences to exclude non-PSS codons, translated them, removed duplicate sequences (sequences remaining: 153), and converted the information into a matrix of five physiochemical descriptor variables that reflect the physical and chemical properties of each amino acid [*44–46*]: z1 (hydrophobicity), z2 (steric bulk), z3 (polarity), z4 and z5 (electronic effects). Using the matrix of z-descriptors, we performed *k*-means clustering with the adegenet’ package in R [*51*], to reveal clusters of sequences likely to have similar functional properties. We then used discriminant analysis of principle components (DAPC) to describe the clusters [*52*].

### 2.6. Runs of homozygosity

Runs of homozygosity (ROH) are stretches of identical haplotypes that occur across homologous chromosomes within the same individual. The length of ROH segments within an individual’s genome depends on whether shared ancestry is recent or ancient; recent inbreeding results in relatively long ROH segments, because recombination has not yet broken up the segments that are identical by descent [*53*]. As mutations accumulate over time, ROH segments break down. We assessed ROH in the Alala genome for three purposes: 1) to estimate the autozygous fraction of the genome fROH [*22*], *i.e.* the total fraction of the genome that is perfectly autozygous (zero heterozygosity); 2) to estimate fROH on a per-contig basis; and 3) to evaluate the effect of allowance for low levels of heterozygosity on estimates of whole-genome fROH and ROH segment lengths. These analyses were conducted using 100 kb sliding windows, with the number of SNPs per sliding window calculated with vcftools [*54*], using the p.vcf file (*i.e.* from the primary contigs) generated during the assembly. A total of four categories of allowable heterozygosity thresholds were tested: ≤ 1, 2, 4, or 10 SNPs per sliding window, which corresponds to 1 SNP per 100kb, 50kb, 25kb or 10kb (*i.e.* ≤ 0.01, ≤ 0.02, ≤ 0.04, and ≤ 0.1 SNPs/kb). Only contigs ≥ 500 kb were included in this analysis, which permits a minimum of five consecutive sliding windows to be included in the ROH segment length estimates. The autozygous fraction of the genome (fROH) was calculated by taking the sum of the number of ROH segments multiplied by the corresponding ROH length and dividing these by the total number of sliding windows considered in the analysis. The ROH lengths were calculated by summing consecutive sliding windows that met criteria of perfect autozygosity or fell within the heterozygosity threshold.

## 3. Results

### 3.1. Genome assembly and quality

The FALCON assembler generated a 1.06 Gbp primary assembly with a contig N50 of 7,737,654 bp across 671 total contigs (Table 1). The diploid assembly process produced 2082 associated haplotype contigs (“haplotigs”) with an estimated length of 0.43 Gb and contig N50 of 455,082 bp (Table 1), implying that about 40% of the genome contained sufficient heterozygosity to be phased into haplotypes by FALCON-Unzip. For comparison, the same assembly process suggest that 75% and 100% of the genomes of two more outbred species, zebra finch and Anna’s hummingbird (*Calypte anna*), contained sufficient heterozygosity to be phased into haplotypes [*26*](Table 1). Compared to other published short-read based avian genomes of similar size, the Alala assembly represents a dramatic decrease in assembly fragmentation, with substantially fewer and longer contigs, and is similar in quality to other long-read *de novo* assemblies [*26*]. The BUSCO analysis indicated that gene completeness was among the highest of any avian genome to date (Figure 1). Collectively, these results suggest that this Alala long-read genome assembly is one of the highest quality avian genomes currently available.

**Table 1.** *De novo* long-read genome assembly statistics comparing PacBio-based primary and secondary haplotypes in three avian species

| Species | PacBio-based primary haplotype | PacBio-based secondary haplotype |
| --- | --- | --- |
| **Alala** |  |  |
| Number of contigs | 671 | 2082 |
| Contig N50 | 7,737,654 bp | 455,082 bp |
| Total size | 1,064,991,496 bp | 432,637,353 bp |
| **Zebra finch <sup>1</sup>** |  |  |
| Number of contigs | 1159 | 2188 |
| Contig N50 | 5,807,022 bp | 2,740,176 bp |
| Total size | 1,138,770,338 bp | 843,915,757 bp |
| **Anna's hummingbird <sup>1</sup>** |  |  |
| Number of contigs | 1076 | 4895 |
| Contig N50 | 5,366,327 bp | 1,073,631 bp |
| Total size | 1,007,374,986 bp | 1,013,746,550 bp |

<sup>1</sup> Korlach, J.; Gedman, G.; Kingan, S. B.; Chin, C.-S.; Howard, J. T.; Audet, J.-N.; Cantin, L. & Jarvis, E. D. *De novo* PacBio long-read and phased avian genome assemblies correct and add to reference genes generated with intermediate and short reads. *GigaScience* **2017** 6, 1-16.

**Figure 1.**
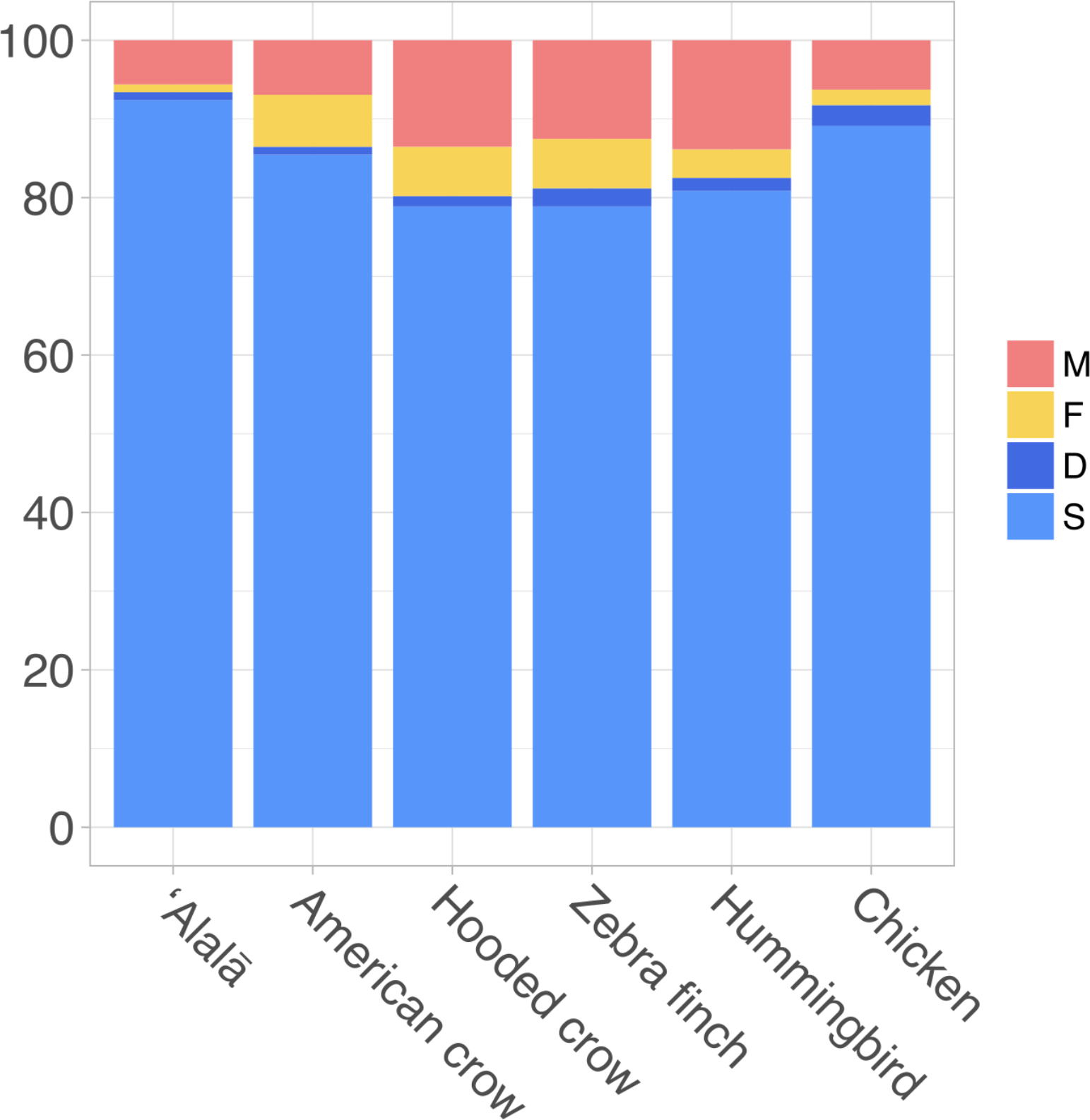
Genome assembly completeness assessed by the recovery of universal single-copy genes (BUSCOs). Percentages refer to complete genes that were found as single (S) or multiple copies (D), as well as fragmented (F) and missing (M) genes. Analyses were based on the BUSCO eukaryote dataset (n = 303 genes).

### 3.2. Mobile and repetitive elements

*De novo* repeat modeling resulted in an Alala-specific repeat library containing 260 families, including 50 LINE (long interspersed element) and 23 LTR (long terminal repeat) families. Only 12% of these had matching entries in Repbase, mostly to endogenous retroviruses (ERVs) and CR1 retrotransposons previously identified in other passerine birds. In addition, several Alala repeat families were partially similar to the large tandem repeat “crowSat1”, a 14 kb satellite that is suspected to be a major heterochromatin component in the hooded crow [*55*]. In contrast, extended matches to Swiss-Prot entries, which might indicate co-opted transposable elements [*56*], were not discovered.RepeatMasker screening identified 10.1% of the Alala assembly as mobile or repetitive sequence, including 3.3% LINEs (exclusively of the CR1 class), 1.1% LTRs (various endogenous retroviruses), and 4.5% unclassified interspersed repeats. The remaining 1.2% was made up of simple repeats and low complexity sequence, including satellites homologous to crowSat1. This estimate did not change noticeably by using a repeat library expanded with avian and ancestral eukaryotic repeats provided by Repbase. The stand-alone analysis of Tandem Repeats Finder revealed 303,030 tandem repeats with a max unit length of 2000 bp, making up 6.9% of the assembly (max repeat size was ~100 kb). This fraction is substantially lower than in the domestic chicken (16.1%), but higher than in the zebra finch (3.7%), Anna’s hummingbird (3.1%), the American crow (2.8%), or the hooded crow (1.8%). However, these results might partially reflect differences in assembly completeness and contiguity, which affect repeat identification (*e.g.* the American crow and hooded crow genome assemblies are short-read based; the Anna’s hummingbird genome was generated and assembled using a similar process as for the Alala). Combined, the RepeatMasker and Tandem Repeats Finder estimates suggest that the Alala assembly contains approximately 15% repetitive DNA. Since large parts of heterochromatic regions routinely elude current-generation sequencing efforts, and repetitive DNA that is sequenced cannot often not be assembled reliably, this is likely an underestimation of the true genome repeat content.

### 3.3. Candidate gene annotations and analysis

We identified the full complement of avian TLR genes in the Alala genome (Table S2), and found them to be highly conserved in number and sequence with respect to the chicken reference [*57*]. Gene structure also proved identical to other birds [*58*], since only a single untranslated 5’ exon seemed to be missing from one prediction. All genes could be classified as functional with full-length reading frames by comparison to other birds. Compared to chicken, notable features included the loss of 140 aa at the 5’ end in *TLR1A*, a ~50 aa indel in *TLR2B*, and a tandem arrangement of two *TLR7* duplicates differing by 16 aa, which we here annotate as *TLR7a* and *TLR7b*. Although a single *TLR7* copy exists in most other birds with annotated TLR repertoires, including zebra and house finch, duplication does occur in several passerine species [*58–62*], indicating that duplicates likely predate the split of the corvid family from other passerines. We did not find evidence that *TLR5* was pseudogenized in Alala, as it is in some passerine species [*63*]; however, based on a single individual this conclusion should be taken with caution.

The MHC class II beta repertoire of the Alala proved to be more complex. We identified seven presumably functional and two ambiguous, but in all likelihood equally functional, genes with open reading frames across all five expected exons (Table S3). This places the Alala within the lower end of the range of known MHC class II B genes relative to other corvids (*e.g.* 7–20 alleles per individual in American crow, jungle crow, and carrion crow [*47*], but note those numbers may include non-functional variants). The Alala genes appear largely conserved compared to zebra finch at the amino acid level, with sequence identity values slightly above 80%. However, it should be noted that this value is derived across entire gene regions, rather than at putative peptide binding regions (in exon 2 for class II B genes), where diversity may be expected to be large. We evaluated exon 2 diversity on a preliminary basis during this supertype analysis, with future work planned to better assess population-level diversity of this region.

The Alala genome also appears to contain a large number of MHC class II B pseudogenes, consistent with expectations for passerines [*e.g. 64*]. In addition to three presumably non-functional genes comprised of exons 2 and 3 (or identifiable parts thereof), we discovered more than 130 sequences homologous to exon 2 on primary contigs, which encodes the highly variable class II histocompatibility antigen beta domain. An additional ~ 30 homologs were found of the more conserved exon 3, which contains the immunoglobulin C1-set domain that mediates T lymphocyte binding (Table S4). Many of these pseudogenes appeared to be highly fragmented, *i.e.* homology could only be established over a short length of 150 bp or less. Sequence identity of these fragments fluctuated widely, ranging from near perfect matches with functional genes to less than 40% at the amino acid level, suggesting a broad age distribution with regard to the time of pseudogenization, and making assembly artefacts inflating the number of pseudogenes unlikely. This might be a reflection of the dynamic evolution of this hyper-variable gene family including repeated expansions and contractions over evolutionary timescales [*65*]. Sequence homology was also visible in adjacent genomic sequence to a lesser degree, suggesting that retrotransposition was not a major mechanism of gene duplication (alignment of 20 randomly chosen pseudogenes ± 250 bp up- and downstream).

Phylogenetically, six of the functional and ambiguous Alala genes comprised a strongly supported clade (Figure S2). These copies only differed by 0–3 amino acids, which were found at or very close to the positively selected sites (PSS) described for other corvids [*47*]. In contrast, only three genes – highly similar copies “a” and “d”, as well as copy “g” – separated to other locations of the phylogenetic tree, albeit with lower support (Figure S2).

### 3.4. MHC functional supertypes

Similarity of the Alala MHC class II B repertoire in terms of functional antigen-binding properties to three other corvids was assessed on the basis of nine PSS [*47*]. Focusing on these PSS in 244 nucleotide sequences, we identified 153 unique amino acid variants. From these, we identified eight functional supertypes (Figure 2), consistent with Eimes *et al.* [*47*]. Congruent with the phylogenetic analysis, only three of the eight supertypes corresponded to Alala, and were also shared by the three other corvid species that were compared (Figure 3). However, it must be noted that sequences from only one Alala were used (compared to 4-6 individuals for each of the other species included in the supertype analysis). No Alala supertypes were discovered to be separate from other corvids.

**Figure 2.**
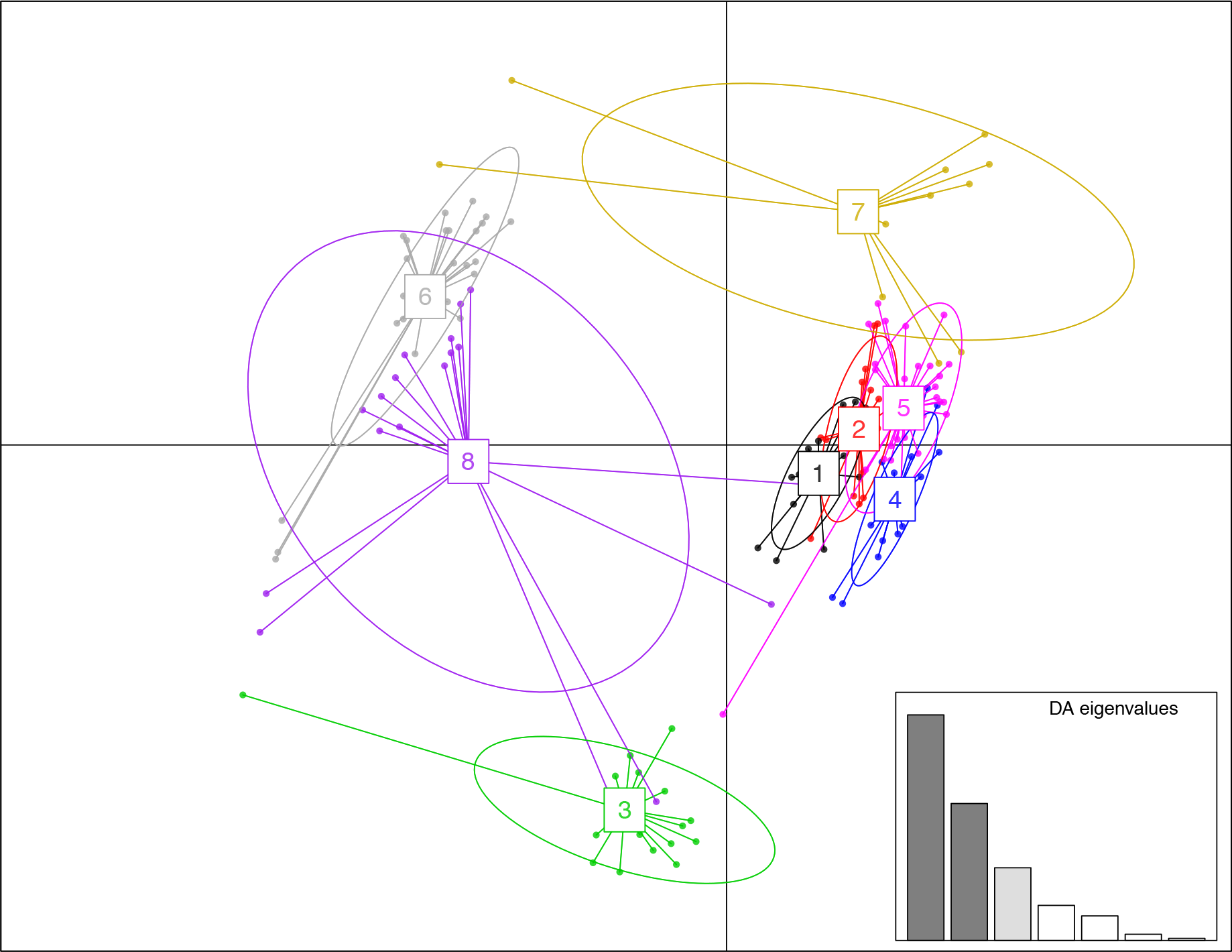
Discriminant Analysis of Principle Components (DAPC) scatterplot of the 8 MHC supertypes. 10 principle components (PC) and three discriminant functions (dimensions) were used to describe the relationship between the clusters. The scatterplot show only the first two discriminant functions (*d* = 2). The bottom graph displays the barplot of eigenvalues for the discriminant analysis. Dark grey, light grey and white bars indicate eigenvalues that were used in the scatterplot, not used in the scatterplot but retained for the analysis, and not retained for the analysis, respectively. Each allele is represented as a dot, and the supertypes as ellipses.

**Figure 3.**
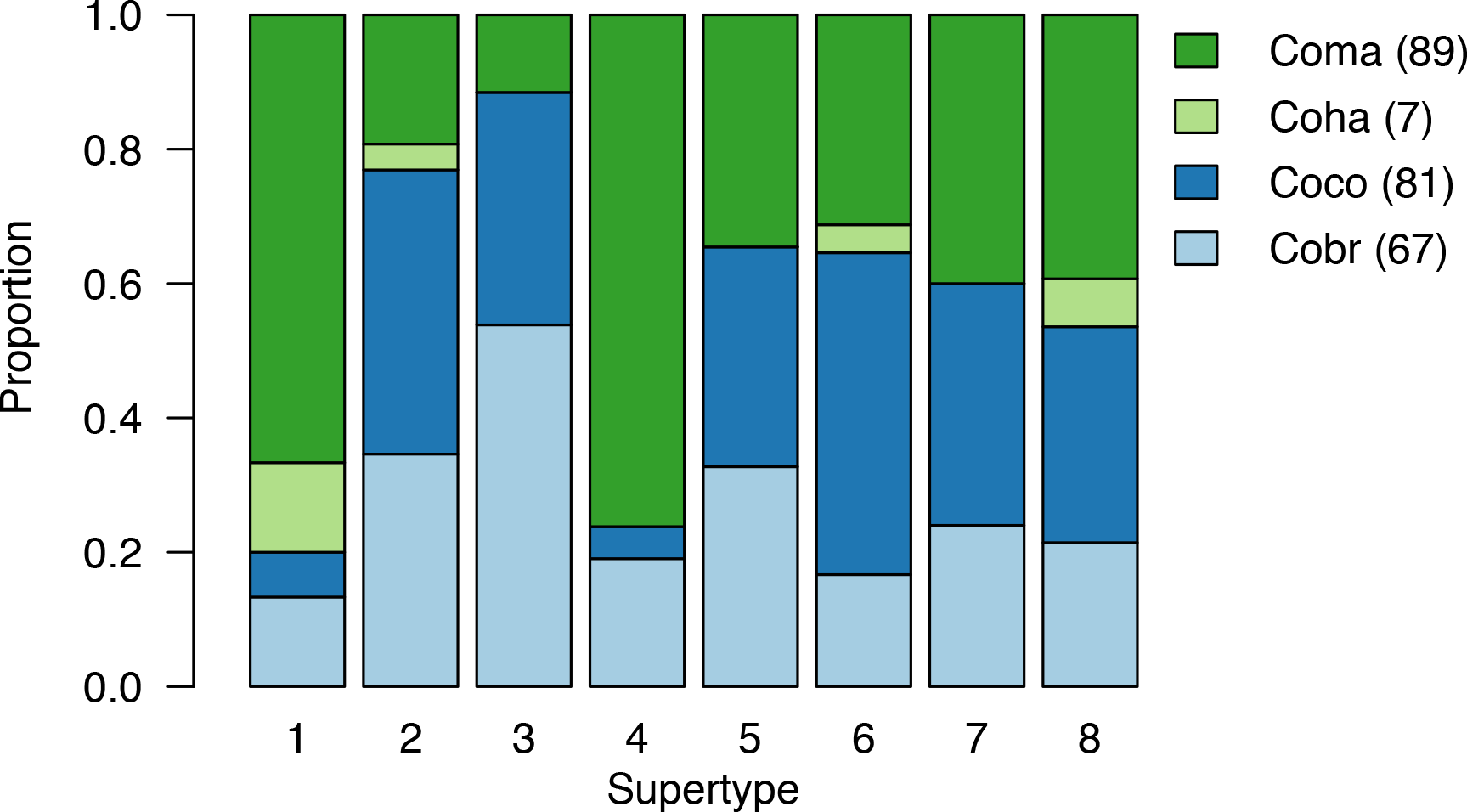
Stacked barplot indicating the representation of each corvid species within each MHC supertype. The three other corvid species were represented across all eight supertypes, while the Alala was represented by three supertypes. Note however that the Alala data were established from a single individual, while the other species’ data represent 4-6 individuals per species [*47*]. In the legend, Coma = jungle crow; Coha = Alala; Coco = carrion crow; Cobr = American crow.

### 3.5. Runs of homozygosity

A total of 413,114 SNPS were detected across the 209 Alala genome contigs that met our minimum ROH analysis requirement of being at least 500 kb in length (1.02 Gb of sequence data, 96% of the genome). Based on the resulting 10,292 sliding windows, the fraction of the genome that was perfectly autozygous (*i.e.*, fROH) was approximately 5.1%; Table S5). The fROH estimates were highly sensitive to allowable levels of heterozygosity. For example, allowing for 1 SNP per 50kb or 25kb in a ROH increased the proportion of whole-genome autozygosity to 28% or 46%, respectively (Table S5). A general trend for low genetic diversity was present across most of the genome, with SNP/kb median and average values of 0.05 and 0.40, respectively. Only a small proportion of sliding windows, 15.6% (1604), were estimated to contain 1 or more SNPs/kb. In terms of individual contigs, the proportion of perfectly autozygous ROHs relative to all sliding windows was highly variable, ranging from a minimum of 0 % to a maximum of 70 % (median 5.3 %; average 1.9 %). The ROH segment lengths were short when restricting ROH criteria to perfectly autozygous, but became increasingly long when allowing or low- to moderate levels of heterozygosity across contigs (Table S5).

## 4. Discussion

Using the PacBio SMRT sequencing technology and FALCON assembly, we generated a high-quality, long-read *de novo* genome assembly for one of the world’s most endangered birds. During the assembly process, FALCON stipulates that if overlapping regions differ by ≥ 5% over extensive distances then the assembler separates the regions into primary and associated (secondary) contigs [*26*]. By definition, regions of the primary assembly that have corresponding associated contigs identify areas in the genome with high degrees of heterozygosity. The genome assembly of a single Alala (studbook #32, Figure S1) highlights the genomic signatures of small population size and inbreeding, because the Alala associated contigs corresponded to a substantially smaller proportion of the genome compared to more outbred species.

### 4.1. Candidate genes

By comparison to many other avian genomes, the Alala genome assembly includes more complete gene sequences than are available for many avian genomes, a crucial factor for annotating complex genomic regions, such as the MHC. The phylogenetically close affiliation and high similarity of six putatively functional Alala MHC class II B genes suggests they may have formed recently, *i.e.* after the split from the other corvids, by multiple rounds of gene duplication. Alternatively, convergent evolution due to strong selection acting on exon 2 (*e.g.* by gene conversion) may also have made these genes more similar to each other and thus mask the gene family’s true evolutionary history [*66,67*]. Genome-scale data of exon 3, or intronic sequence from other corvids and additional Alala individuals may be required to resolve this question. Three Alala MHC class II B sequences did not cluster together. While a lack of support along the backbone of the phylogenetic tree prevented the identification of clear orthologs with other species, these three sequences may represent remnants of evolutionarily distinct gene lineages. Multiple MHC class II B lineages have been identified in other corvids, suggesting that the gene family expanded prior to the radiation of the corvid family [*40,47*]. It should be noted that no Alala genes were found to fall into any of several MHC class II B groups consisting of genes from all or almost all other corvids investigated here (Figure S2). It is thus possible that such Alala genes were lost in the evolutionary lineage leading to the Alala, or even very recently in the species’ population history. This hypothesis is supported by the high number of gene fragments, and high sequence similarity between copy “g” and pseudogene “p1”, suggesting at least one more recent pseudogenization event. However, these observations are based on only a single individual and should be interpreted with caution. Genome data from additional specimens would be required to gauge within-species diversity and distribution of different MHC class II B lineages, and put these data into the context of corvid and Alala evolution. Additionally, our functional supertype analysis suggests that while nucleotide sequences may differ, between Alala and other corvids, much similarity exists when it comes to pathogen-binding properties.

### 4.2. Runs of homozygosity

The SNP encounter rate estimate in the Alala is about 1 SNP per 2,477 bp (413,114 SNPs identified from 1.02 Gb of sequence data). This value is considerably lower than empirical estimates obtained from genomes of other avian species, for example, 1 SNP per 330 bp in *Ficedula* flycatchers [*68*], 1 SNP per 256 bp in Hawaii Amakihi (*Hemignathus virens* [*69*]) and 1 SNP per 935 bp in turkey (*Meleagris* spp. [*70*]). The Anna’s hummingbird genome [*26*] revealed 1,841,030 variants (*i.e.* 1 SNP per 501 bp) across n = 283 contigs of minimum length of 500 kb (923.1 Mb). While caution should be used when making comparisons between genomes that differ by sequencing technologies, genome assembly pipelines, and other computational settings (addressed in more detail below), the paucity of SNPs in the Alala genome is not surprising because of the overall low population size of Alala and this particular bird’s high pedigree inbreeding coefficient (0.25). In the examples noted here, the Alala genome was generated and assembled in a similar fashion to that of the Anna’s hummingbird, the latter of which had a SNP encounter rate almost five times as frequent. Certainly, the presence of chromosomes showing very low heterozygosity in the Alala is consistent with empirical observations made of ROHs in turkey [*70*], and large stretches of very low heterozygosity in Hawaii Amakihi [*69*]. The contrast between highly variable sliding windows and regions with modest variability suggests that the PacBio assembly pipeline used here is sensitive to calling SNPs across a range of heterozygosities, and that low diversity observed for this genome is not solely an artifact of the assembly pipeline.

The cumulative lengths of ROH are an indication of shared ancestry and can be used to gauge whether inbreeding events occurred within recent or distant generations [*53*]. Recombination events break long tracts into smaller pieces, thus numerous short tracts are consistent with distant shared ancestry. In this Alala genome the majority of tracts of ROHs were short, even when allowing for 1 SNP per 100 kb (Table S5). Yet, if allowable heterozygosity is increased to 1 SNP per 25,000 bp then 9.8% (104 of 1054) of all tracts would exceed 1 Mb. Given this measure’s high sensitivity to filtering criteria interpretation of results would be aided considerably by whole genome sequencing of additional progeny or parents. The rate of sequencing error will also affect the estimations of homozygosity and, correspondingly, the lengths of runs of homozygosity.

Several factors confound comparability of ROH across studies and taxa. These include: lack of consensus definition for ROH; differences in sequencing platforms and associated sequencing errors; the variant-calling pipeline; and computational settings [*e.g. 71–73*]. The ROH estimates in this study are drawn from a single genome with high depth of sequencing. In contrast, measures of ROH can be obtained from high-density SNP arrays by quantifying the length of rows of homozygous SNPs relative to a reference, the results of which are sensitive to SNP chip density, and may miss unmeasured variants between the markers [*71,72*]. The density of SNPs across the genome also impacts the minimum length of detection for a ROH, for example, reliable detected of ROHs as short as 100 kb versus 510 Mb using a less dense panel of markers [*53*]. Comparability of ROH results between avian and mammalian study species are further diminished by biological variability in chromosome lengths. Birds, with numerous microchromosomes, are expected to have shorter ROH than mammals, simply because of differences in chromosome lengths.

### 4.3. Applications to Alala conservation

The Alala genome assembly resulting from long-read sequencing data provides a high-quality reference genome that is enabling downstream comparative, population, and conservation applications. Prior to this study, molecular work using limited genetic markers identified low diversity in the species [*20,21*], and pedigree analysis identified inbreeding effects on hatching success, as well as skewed founder representation [*15,18*]. Based on these studies, we identified a need to generate genomic resources to aid conservation management. These included a more detailed picture of population-level genomic diversity and genetic load, as well as more accurate estimates of molecular relatedness to assist with choosing breeding pairs. Genomic resources were also desired to begin investigations of the basis of traits such as poor hatching success, as ~ 61% of fertile Alala eggs fail [*15*], compared with ~ 10% in most birds [*74*]. Since the generation of this Alala assembly, several projects have been initiated that rely heavily on use of this new resource. For example, we are currently using targeted SNP-capture to compare genomic diversity in museum and modern Alala, to better understand the impact of population bottlenecks over the past 100 years, and to provide a clearer picture of how much diversity can likely be maintained into the future (manuscript in preparation). So far, these analyses have benefitted from our high-quality genome assembly by allowing identification and removal of sequence reads that map to multiple genomic locations, and through testing and control for linkage disequilibrium (*i.e.* better quality filters). More recently, we have begun reduced representation library sequencing using restriction site-associated DNA sequencing (RAD-seq) strategies to genotype every individual Alala, so that we can inform the choice of breeding pairs in captivity by using realized relatedness estimates that likely correlate better with phenotypic trait information compared to pedigree-based relatedness estimates [*e.g. 11*]. Analytic methods for RADseq data show less bias when a suitable reference genome is included [*75*]. We have also initiated population-level assessments of candidate genes, which were aided by our annotations. Long-term, we plan to establish whole-genome data for multiple individuals, and to incorporate all of these datasets into models that test for genomic basis of particular phenotypic traits (*e.g.* poor hatching success), as well as models of mate choice and other behaviors.

Genomic data derived from our analyses are an essential component of the current and future recovery of the Alala. Although pair selection and managed breeding using the pedigree has kept the inbreeding level of the Alala population at a relatively low level over the past 20 years [*18*], the intensive and ongoing conservation management of the species requires a more detailed approach [*e.g. 11*]. In conjunction with ongoing conservation breeding efforts, a comprehensive reintroduction program is underway in an effort to re-establish this formerly extinct-in-the-wild species into its native forest habitat. Early indications of the reintroduction effort are promising, with a small founder population surviving in the wild at the time of writing. Ongoing management decisions for the breeding and release of particular individuals will have implications for the recovery of the species. As the size of both the captive and wild Alala populations continue to increase, the integration of genomic data as part of the conservation management effort will help to maximize the genetic health of the species well into the future.

## Supplementary Materials

The following are available online at www.mdpi.com/xxx/s1, Tables S1–S5, Figures S1–S2. Primary and secondary genome assemblies will be available from Genbank, Accessions: (to be provided during review).

## Ethics Approval

All procedures on live animals were approved by the Institutional Animal Care and Use Committee (IACUC) of San Diego Zoo Global (15-012 and 16-009).

## Author Contributions

Conceptualization, Jolene T. Sutton, Cynthia C. Steiner, Jenny Gu, Bryce M. Masuda and Oliver Ryder; Data curation, Richard Hall, Primo Baybayan and Jill Muehling; Formal analysis, Jolene T. Sutton, Martin Helmkampf, Cynthia C. Steiner, M. Renee Bellinger, Jonas Korlach and Sarah Kingan; Methodology, Jonas Korlach, Richard Hall, Primo Baybayan and Jill Muehling; Writing – original draft, Jolene T. Sutton, Martin Helmkampf, Cynthia C. Steiner, M. Renee Bellinger, Jonas Korlach, Bryce M. Masuda and Oliver Ryder.

## Funding

Funding for the conservation breeding and reintroduction efforts has been provided by the San Diego Zoo Global, U.S. Fish and Wildlife Service, Hawaii Division of Forestry and Wildlife, National Fish and Wildlife Foundation, the Max and Yetta Karasik Foundation, the Moore Family Foundation, American Forests, and numerous anonymous donors. The material presented here is partially based upon work supported by the National Science Foundation under Grant No. 1345247. Any opinions, findings, and conclusions or recommendations expressed in this material are those of the author(s) and do not necessarily reflect the views of the National Science Foundation.

## Acknowledgments

Many thanks to the hard working staff, interns, and volunteers who care for and propagate Alala at the facilities on Maui (Maui Bird Conservation Center) and Hawaii (Keauhou Bird Conservation Center).

## Conflicts of Interest

JK, RH, PB, JM, JG, and SK are full-time employees at Pacific Biosciences, a company developing single-molecule sequencing technologies. All other authors declare that they have no competing financial interests

---

**Table S1.** Accessions of references used to annotate candidate immunity genes.

| Gene group | Reference species | Reference accession | Source |
| --- | --- | --- | --- |
| MHC | zebra finch | UniProtKB: H1A2M7_TAEGU | 29 |
|  |  | UniProtKB: H0ZVH6_TAEGU | 29 |
|  |  | UniProtKB: H0ZU54_TAEGU | 29 |
|  |  | UniProtKB: H1A3G3_TAEGU | 29 |
|  |  | UniProtKB: H0ZXA5_TAEGU | 29 |
|  |  | UniProtKB: H0ZZB1_TAEGU | 29 |
|  |  | UniProtKB: H1A0J1_TAEGU | 29 |
|  |  | UniProtKB: H0ZWA2_TAEGU | 29 |
|  |  | UniProtKB: H0ZWI5_TAEGU | 29 |
|  |  | UniProtKB: H0ZWL2_TAEGU | 29 |
|  |  | UniProtKB: H0ZXL7_TAEGU | 29 |
| TLR | chicken | GenBank: NM_001007488.4 | 76 |
|  |  | GenBank: NM_001081709.3 | 77 |
|  |  | GenBank: NM_204278.1 | 78 |
|  |  | GenBank: NM_001161650.1 | 79 |
|  |  | GenBank: NM_001011691.3 | 80 |
|  |  | GenBank: NM_001030693.1 | 81 |
|  |  | GenBank: NM_001024586.1 | 82 |
|  |  | GenBank: NM_001011688.2 | 83 |
|  |  | GenBank: NM_001037835.1 | 84 |
|  |  | GenBank: NM_001030558.1 | 81 |

**Table S2.**
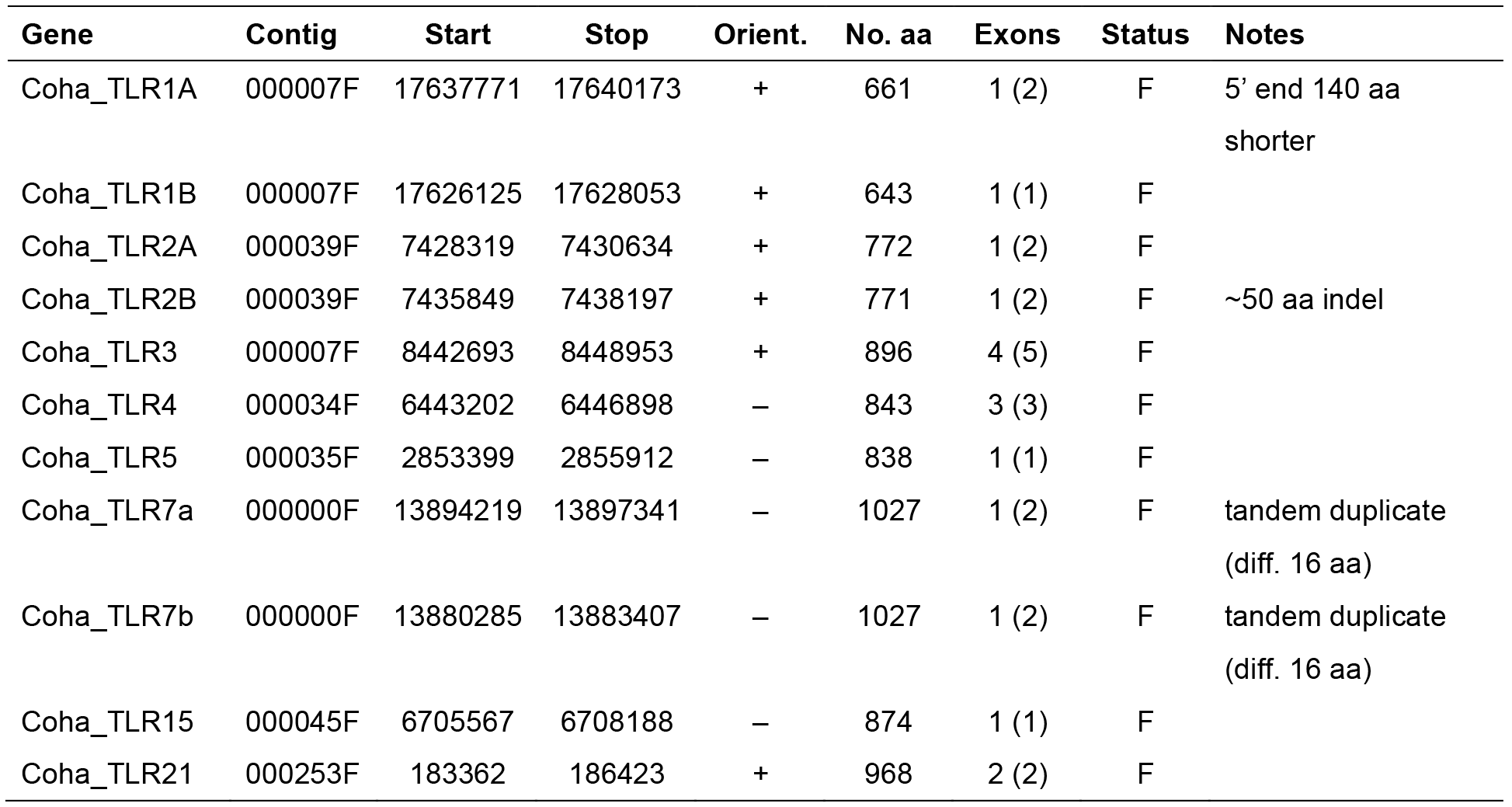
Alala toll-like receptor (TLR) genes, following the nomenclature suggested by Temperley et al. (2008). Start and stop refer to the position of the start and stop codons on the contig. Exons indicates the number of exons identified in the absence of transcriptional evidence, with the number of exons in the *G. gallus* reference given in parentheses. All predicted genes appear to be complete, suggesting they are functional (F). Notes are provided with respect to the *G. gallus* reference, where applicable.

| Gene | Contig | Start | Stop | Orient. | No. aa | Exons | Status | Notes |
| --- | --- | --- | --- | --- | --- | --- | --- | --- |
| Coha_TLR1A | 000007F | 17637771 | 17640173 | + | 661 | 1 (2) | F | 5' end 140 aa shorter |
| Coha_TLR1B | 000007F | 17626125 | 17628053 | + | 643 | 1 (1) | F |  |
| Coha_TLR2A | 000039F | 7428319 | 7430634 | + | 772 | 1 (2) | F |  |
| Coha_TLR2B | 000039F | 7435849 | 7438197 | + | 771 | 1 (2) | F | ~50 aa indel |
| Coha_TLR3 | 000007F | 8442693 | 8448953 | + | 896 | 4 (5) | F |  |
| Coha_TLR4 | 000034F | 6443202 | 6446898 | – | 843 | 3 (3) | F |  |
| Coha_TLR5 | 000035F | 2853399 | 2855912 | – | 838 | 1 (1) | F |  |
| Coha_TLR7a | 000000F | 13894219 | 13897341 | – | 1027 | 1 (2) | F | tandem duplicate (diff. 16 aa) |
| Coha_TLR7b | 000000F | 13880285 | 13883407 | – | 1027 | 1 (2) | F | tandem duplicate (diff. 16 aa) |
| Coha_TLR15 | 000045F | 6705567 | 6708188 | – | 874 | 1 (1) | F |  |
| Coha_TLR21 | 000253F | 183362 | 186423 | + | 968 | 2 (2) | F |  |

**Table S3.** Alala MHC class II beta genes. Beginning and end coordinates refer to initial, uncurated gene models, and may or may not coincide with start and stop codons. The Exons column provides the number of exons identified in the absence of transcriptional evidence. The Status column indicates whether the predicted gene is presumably functional (F), ambiguous (A) or a pseudogene (P) according to the definition given in the main text. Only pseudogenes containing both exon 2 and 3 are listed. “Frameshift fixed” denotes gene models that were fully recovered by the insertion of a single nucleotide at a homopolymer (suggesting a sequencing error).

| Gene | Contig | Begin | End | Orient. | No. aa | Exons | Status | Notes |
| --- | --- | --- | --- | --- | --- | --- | --- | --- |
| Coha_MHCIIB_a | 000257F | 17839 | 19521 | – | 250 | 5 | F |  |
| Coha_MHCIIB_b | 000357F | 54501 | 55649 | + | 250 | 5 | F |  |
| Coha_MHCIIB_c | 000357F | 69097 | 70245 | + | 250 | 5 | F |  |
| Coha_MHCIIB_d | 000357F | 80536 | 81696 | + | 250 | 5 | F |  |
| Coha_MHCIIB_e | 000485F_001 | 9214 | 10361 | + | 250 | 5 | A | Frameshift fixed |
| Coha_MHCIIB_f | 000485F_001 | 29737 | 30884 | + | 250 | 5 | F |  |
| Coha_MHCIIB_g | 000485F_001 | 46119 | 47225 | + | 250 | 5 | F |  |
| Coha_MHCIIB_h | 000868F | 15490 | 16637 | + | 250 | 5 | F |  |
| Coha_MHCIIB_i | 000868F | 26421 | 27564 | + | 249 | 5 | A | Frameshift fixed |
| Coha_MHCIIB_p1 | 000868F | 4152 | 5188 | + | 251 | 5 | P | Divergent, multiple frameshifts, non-canonical splice site |
| Coha_MHCIIB_p2 | 000251F | 76819 | 77558 | + | 212 | 3 | P | Fragmentary (exons 1–3), frameshift, mutated splice site |
| Coha_MHCIIB_p3 | 000219F | 136202 | 137011 | + | 99 | 3 | P | Fragmentary (exons 1–3), multiple frameshifts |

**Table S4.**
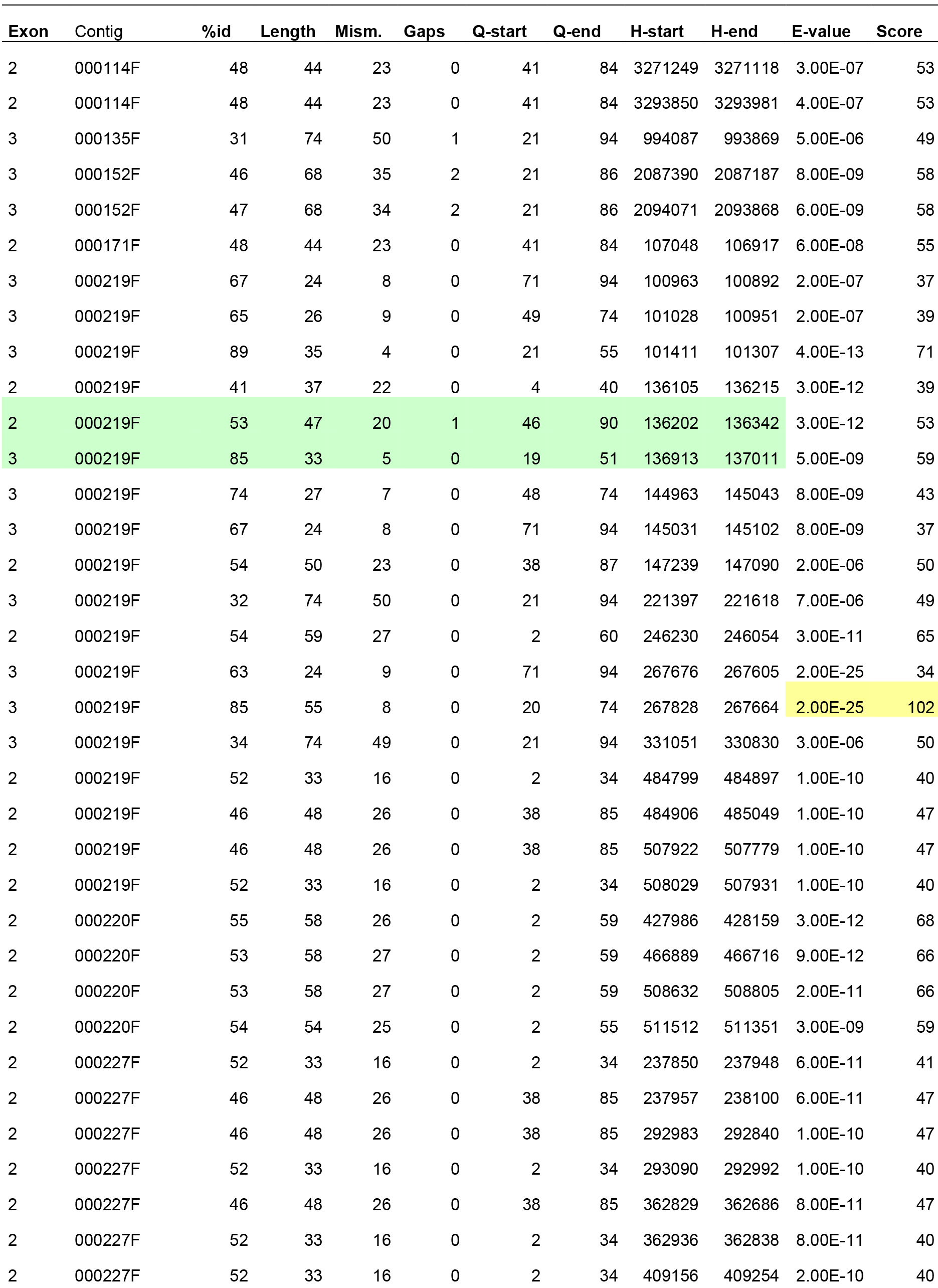
Homologs of MHC class II B exons 2 and 3 in the Alala genome. Identity (%id), mismatches (Mism.), gaps, query start and end positions (Q-start, Q-end), hit start and end positions (H-start, H-end), e-value and score refer to Blast results using Coha_MHCIIB_b exons 2 and 3 as query. Bit scores > 100 are highlighted in yellow. Consecutive hits on the same contig within 1500 bp (in green blocks) were annotated further as MHC class II B candidate genes (see Table S2). The remaining hits are fragments that may represent ancient, pseudogenized copies.

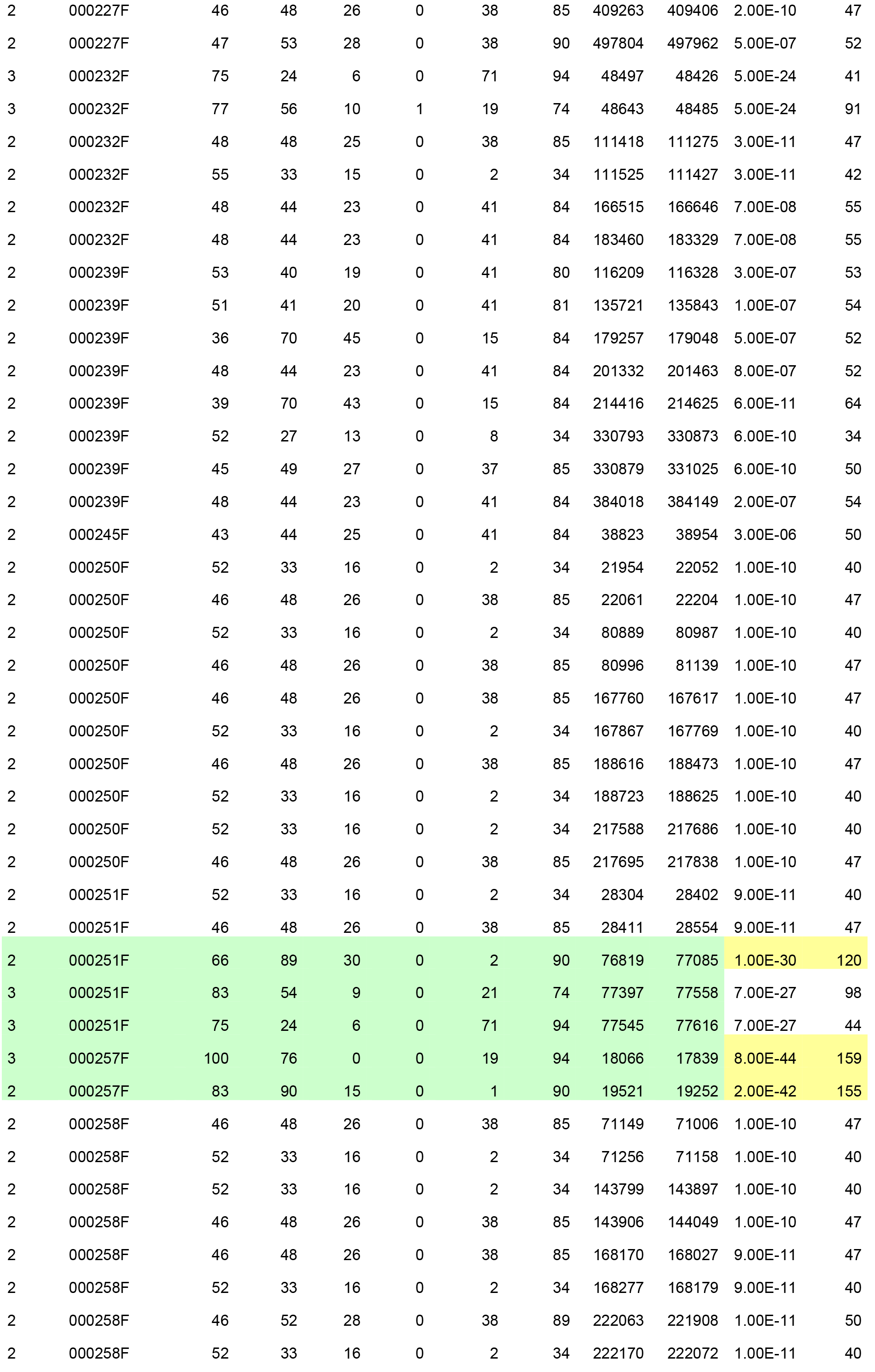

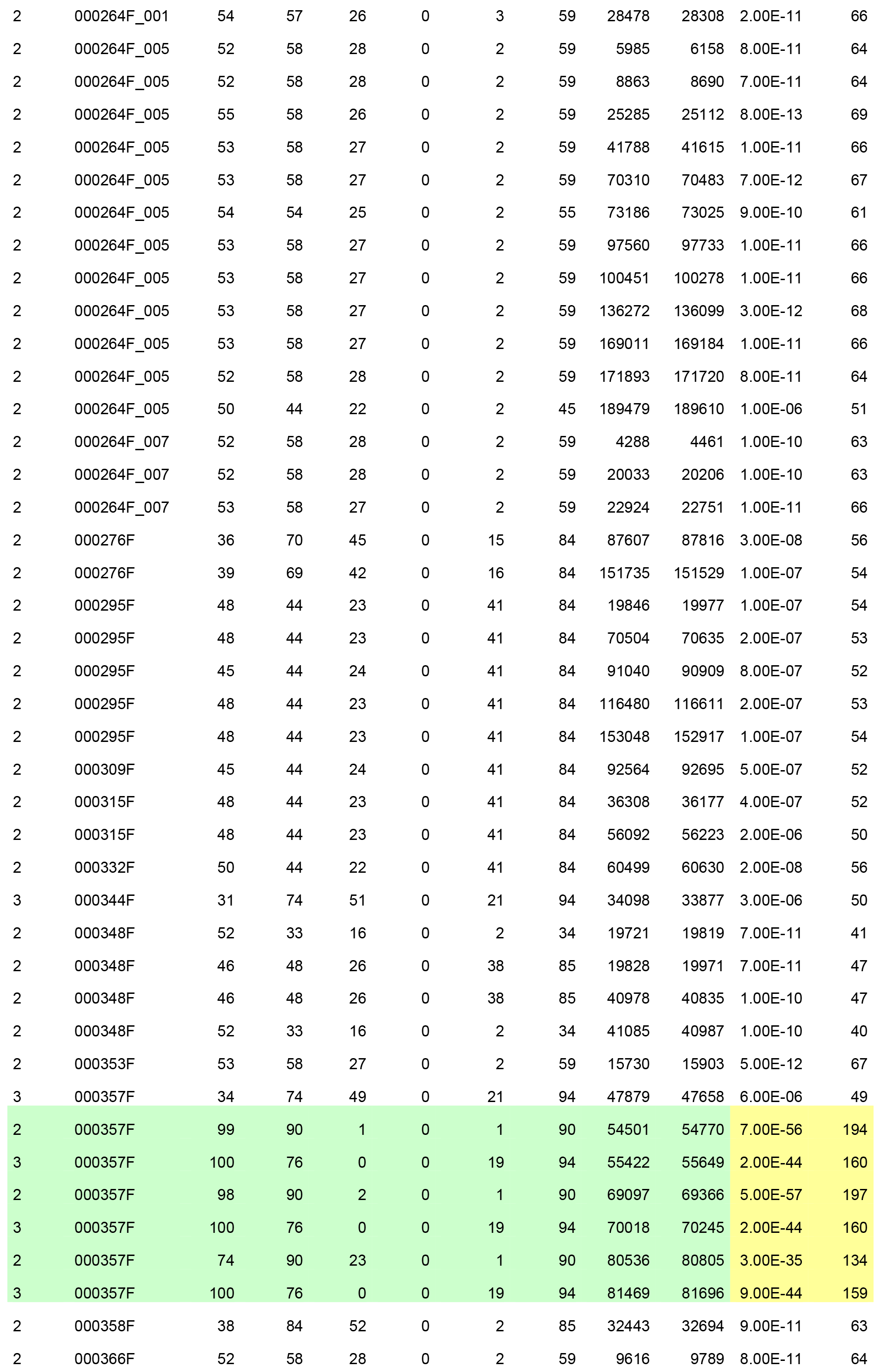

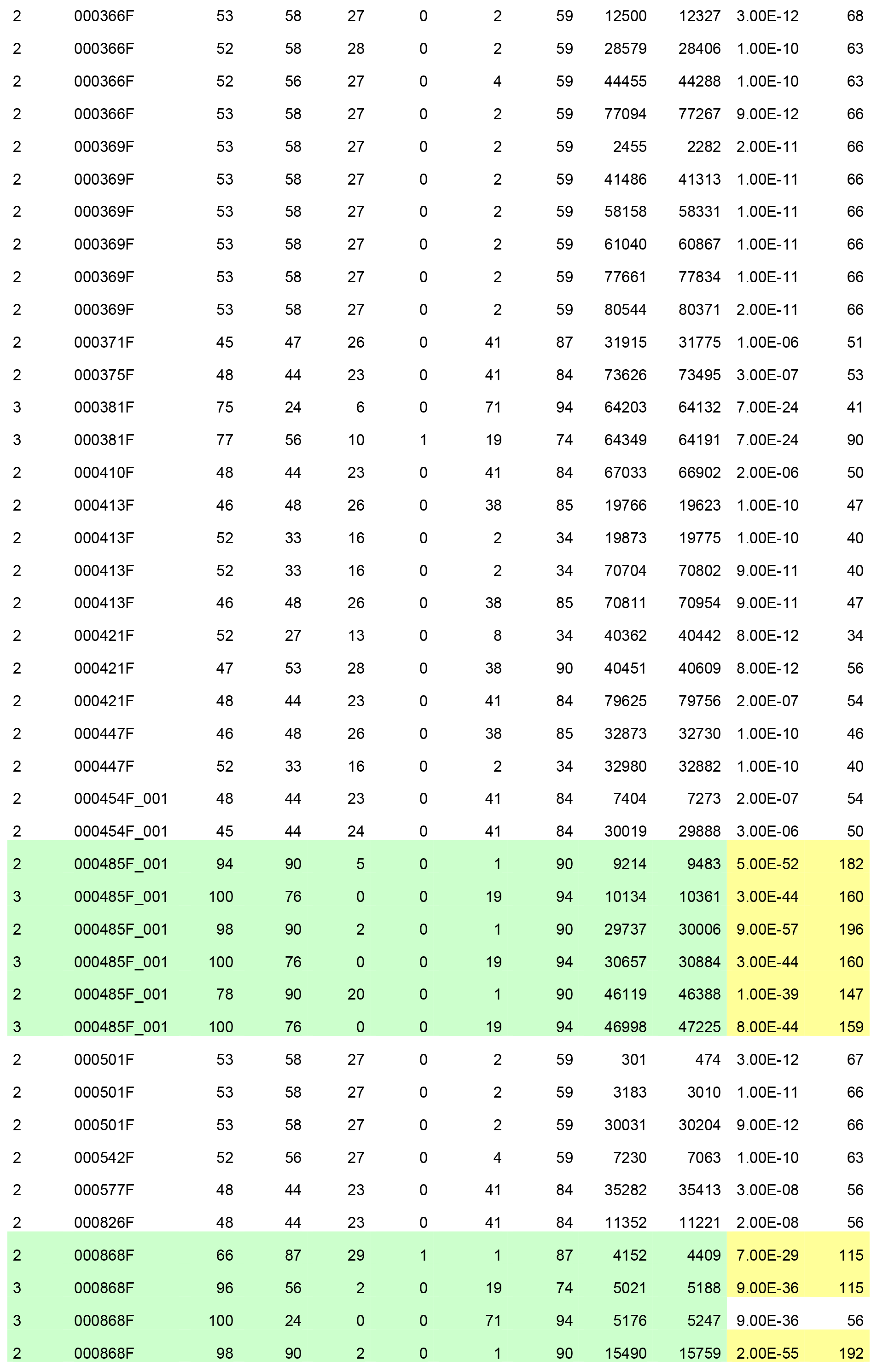

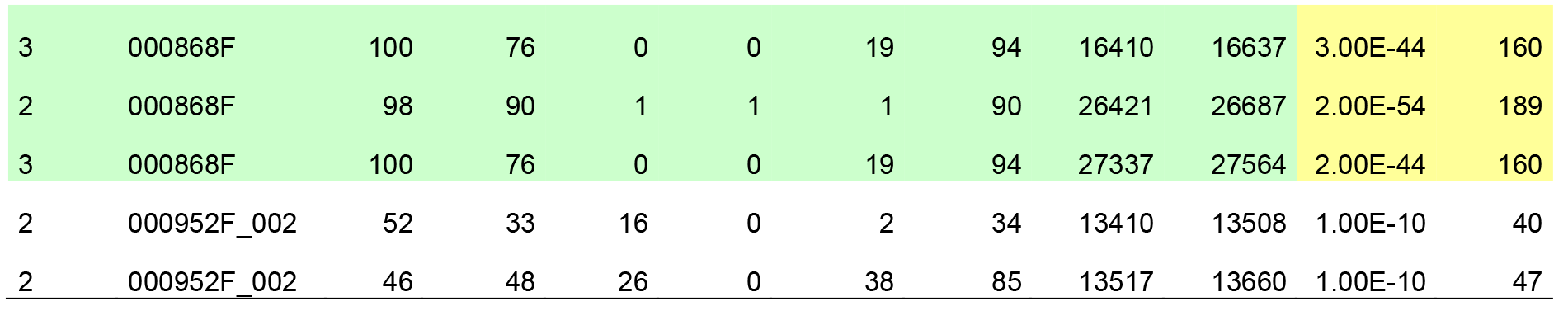

**Table S5.**
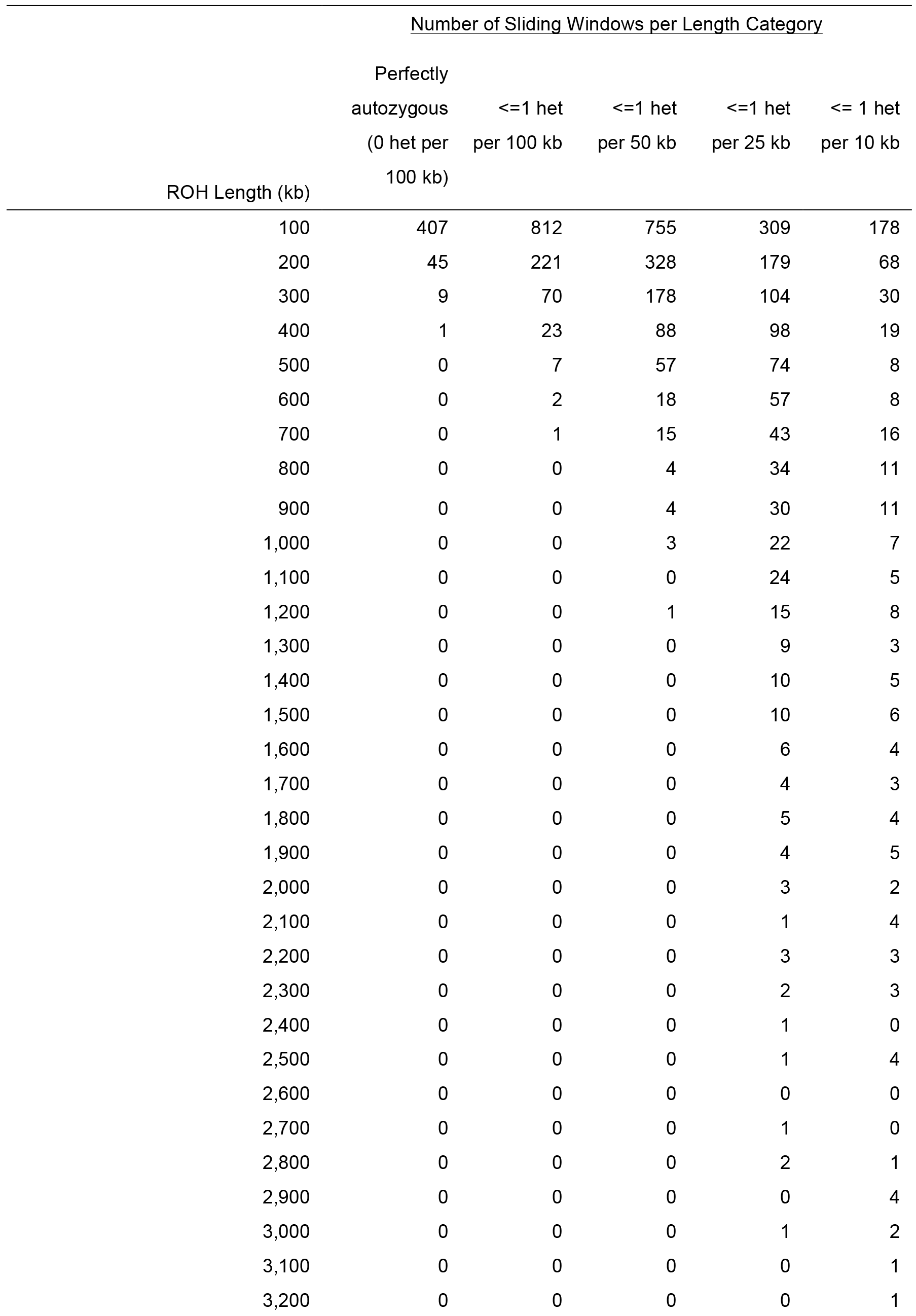
Analysis of runs of homozygosity across 209 Alala genome contigs with lengths ≥ 500kb. Each row indicates the number ROH segments observed per ROH length category. Data are provided under conditions of complete autozygosity and four heterozygosity thresholds.The autozygous fraction of the genome (fROH) was calculated by taking the sum of the number of ROH segments multiplied by the corresponding ROH length. The ROH segments are based on 100kb sliding window intervals.

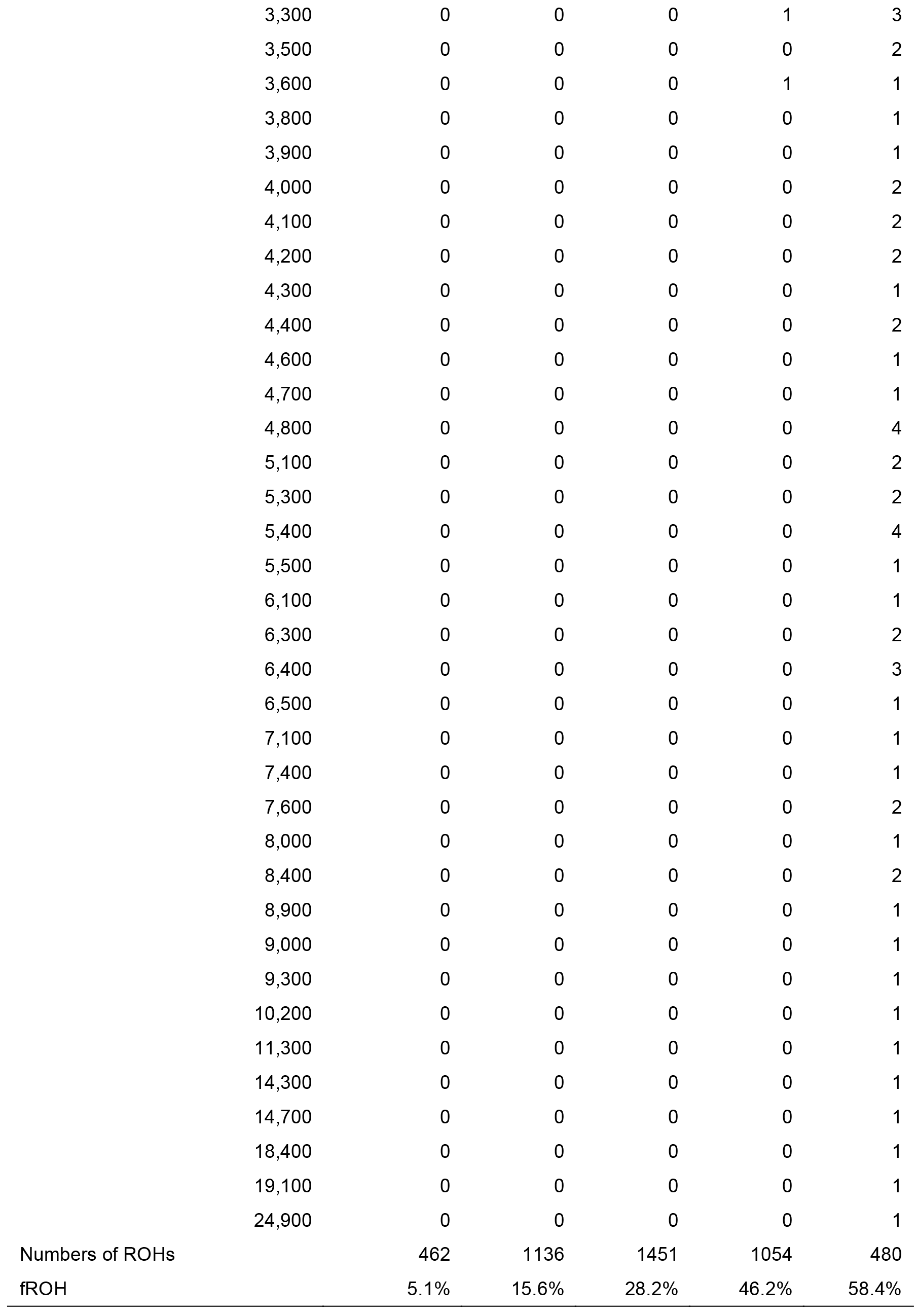

**Figure S1.**
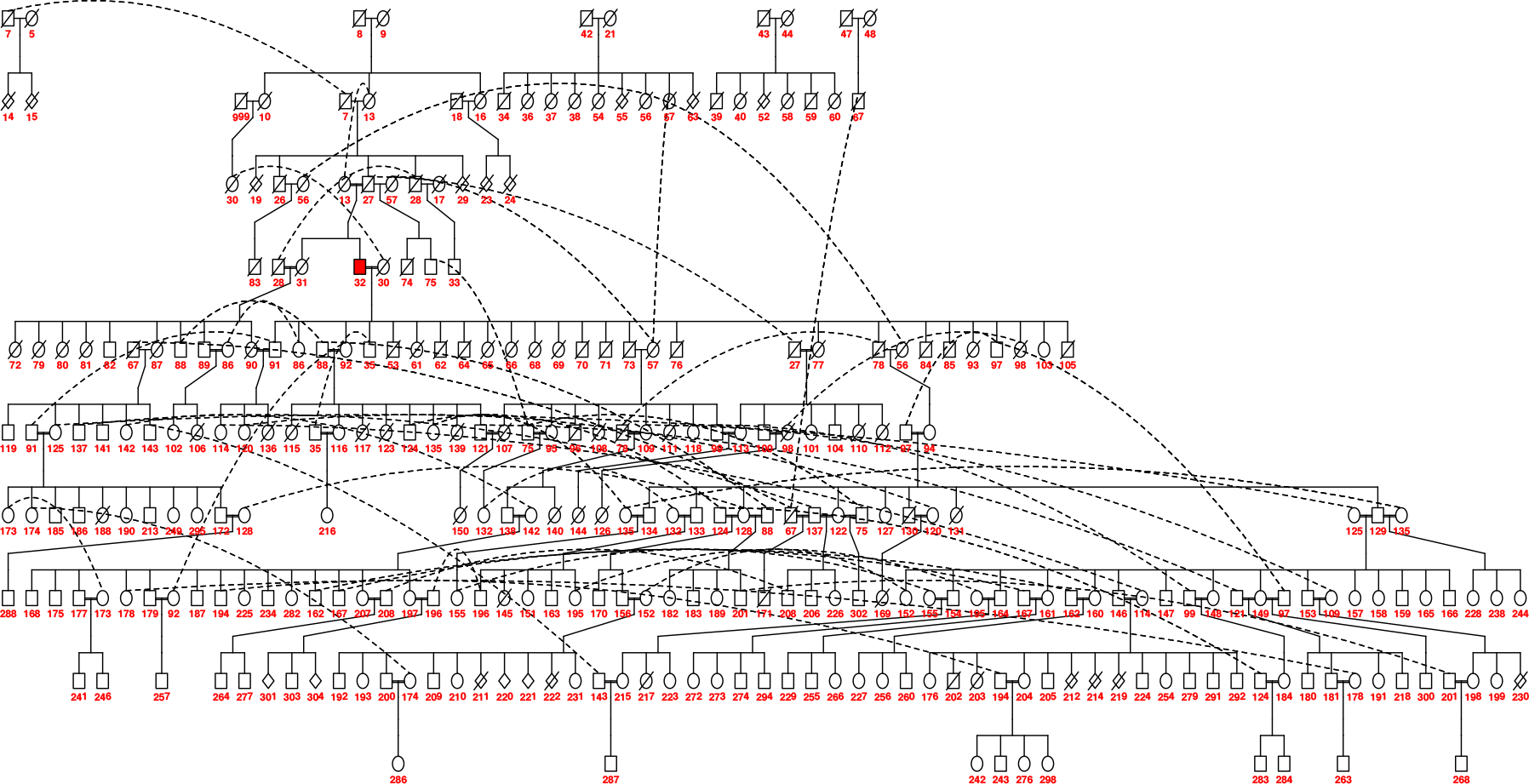
Alala pedigree. The sequenced individual, studbook 32 (named Hōike i ka pono) is shaded. Dashed lines indicate individuals that are represented in multiple positions in the pedigree (*e.g*. overlapping generations).

**Figure S2.**
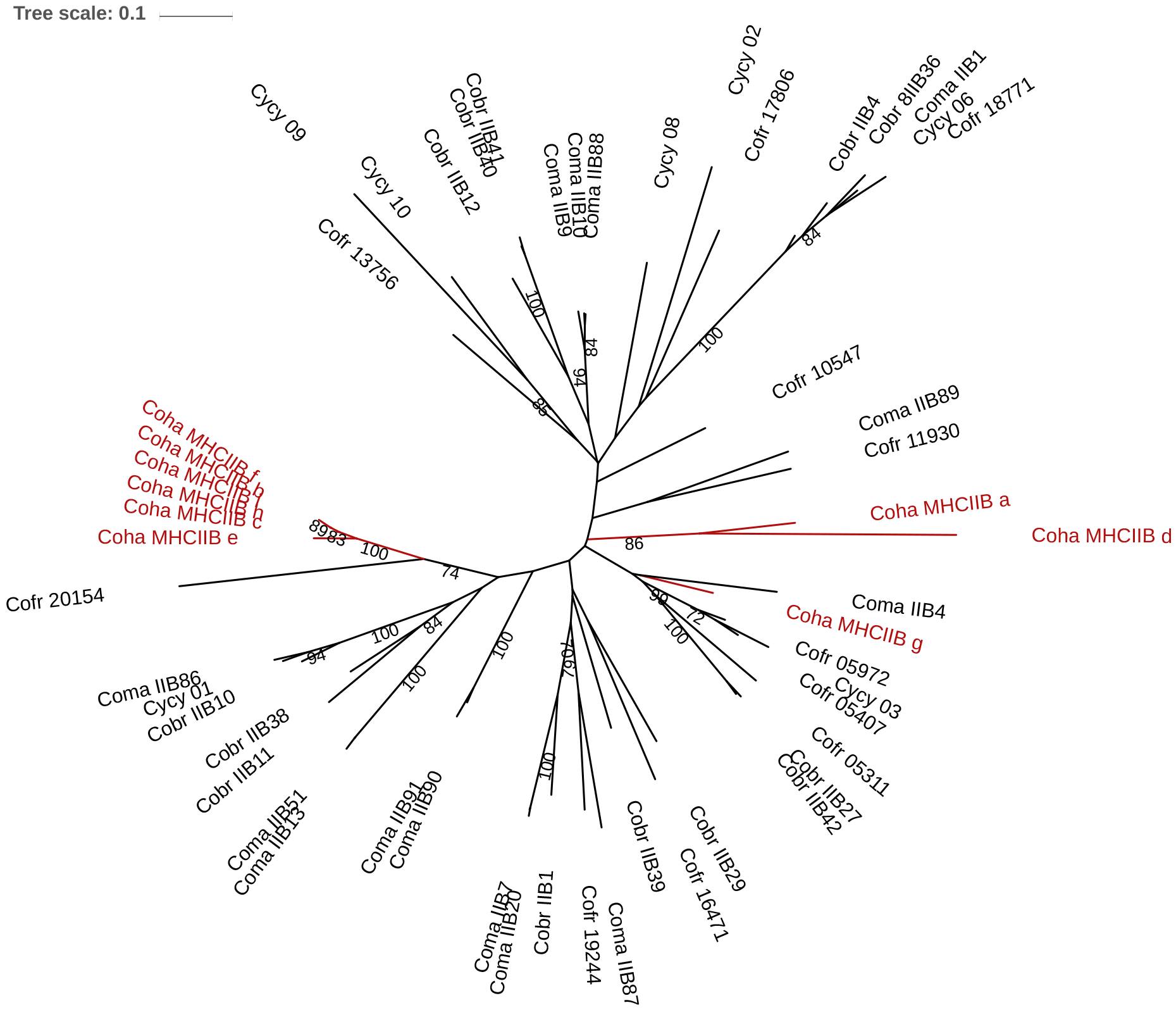
Unrooted maximum likelihood tree of corvid MHC class II beta genes, exon 2. Functional genes predicted from the Alala assembly are highlighted in red (prefix Coha). Other species represented include *C. brachyrhynchos* (American crow, Cobr), *C. macrorhynchos* (jungle crow, Coma), *C. frugilegus* (Asian rook, Cofr) and *Cyanopica cyanus* (azure-winged magpie, Cycy). All genes were obtained from one individual per species (Eimes et al. 2016 and pers. comm.). Confidence values are given for nodes with bootstrap support > 70%.

